# Segmentation-guided photon pooling enables robust single cell analysis and fast fluorescence lifetime imaging microscopy

**DOI:** 10.1101/2025.09.30.679660

**Authors:** Kayvan Samimi, Danielle E. Desa, Xiaotian Zhang, Dan L. Pham, Rupsa Datta, Melissa C. Skala

## Abstract

Fluorescence lifetime imaging microscopy (FLIM) can probe the metabolic environment of living cells in a label-free and non-invasive manner. However, endogenous fluorophores have low absorption and quantum yields, which necessitates long integration times to acquire the high photon counts needed for accurate pixel-wise multi-exponential decay fitting. Here, we present a ‘region-of-interest’ photon pooling technique to expedite label-free, single cell FLIM acquisition and analysis. As a result, we achieved single-cell metabolic information at intervals as low as one second and acquired large FLIM mosaics 15 times faster than would be possible with conventional pixel-level analysis. This technique is computationally light, does not require machine learning algorithms, and has been integrated with commonly used analysis software and file types.

## 1. Introduction

Fluorescence lifetime imaging microscopy (FLIM) has become a powerful tool for assessing living biological specimens. FLIM is sensitive to the local environment of the fluorophore, including pH, oxygen concentration, temperature, and conformational changes with protein binding.^1,2^ Autofluorescent metabolic coenzymes, reduced nicotinamide adenine dinucleotide (phosphate) (NAD(P)H) and oxidized flavin adenine dinucleotide (FAD), provide a label-free source of contrast for FLIM.^3^ Therefore, FLIM of NAD(P)H and FAD can be used to study changes in cell metabolism over time in a nondestructive manner.^4,5^

However, measuring lifetime decays from autofluorescent biomolecules presents unique challenges. First, the quantum yield, defined as the ratio of emitted to absorbed photons, of autofluorescent biomolecules (*e*.*g*., NAD(P)H) is 30–40× lower than conventional engineered fluorophores (*e*.*g*., green fluorescent protein, tdTomato).^6–9^ Second, resolving the fluorescence lifetime of autofluorescent biomolecules requires more complex analysis because these molecules exist in multiple binding states. For example, NAD(P)H can exist freely in a cell or bind to over 300 other molecules in various metabolic and biosynthetic reactions.^10^ NAD(P)H can also self-quench when unbound, resulting in a shorter lifetime (∼400 ps) relative to its bound state (∼2–5 ns).^9,11^ More photons (>1000/pixel after spatial binning)^12^ are therefore needed for multiexponential fitting to accurately recover complex autofluorescence lifetimes, often resulting in longer image acquisition times and potential photodamage. Acquisition times on the order of one minute per autofluorescence FLIM image are common. This restricts the ability of autofluorescence FLIM to capture highly dynamic (∼1 second) biological processes in live cells.^13^

FLIM hardware solutions provide one approach to reduce autofluorescence FLIM acquisition times. Time-correlated single photon counting (TCSPC) is the most sensitive and photon-efficient time-resolved acquisition technique in low-light settings, providing excellent signal-to-noise ratio (SNR) and timing resolution.^13,14^ While most TCSPC FLIM implementations rely on single-point laser scanning systems, parallel excitation can be used to simultaneously illuminate multiple points in the sample, and thereby reduce acquisition times.^3,13^ For example, multifocal FLIM^15,16^ or light sheet illumination^17,18^ can be combined with high-speed detectors such as single photon avalanche photodiode (SPAD) arrays^19–21^ to produce sufficient SNR to resolve TCSPC lifetime decays using faster acquisition times than single-point laser scanning methods^15–18^.

Alternatively, post-processing techniques can be used to compensate for faster acquisition times and the resulting low SNR FLIM images. Lifetime decays are typically fit in a pixel-wise manner using iterative re-convolution with the instrument response function (IRF) through weighted least squares (WLS) or maximum-likelihood estimation (MLE) optimization, which require hundreds of photons per pixel to achieve accurate fits.^3,13^ Local spatial pixel binning (*i*.*e*., aggregation of neighboring pixel decays in a 3×3, 5×5, or larger kernel) is regularly employed in pixel-wise fitting (*e*.*g*., SPCImage software) to achieve the required photon counts. Alternative approaches such as global fitting algorithms, which assign ‘best guesses’ to a subset of the lifetime fit parameters based on spatial distributions within a sample, can decrease the number of photons per pixel needed for fitting.^22–24^ Similarly, Bayesian analysis determines the likelihood function and lifetime from a prior fluorescence decay to establish and then maximize the posterior distribution of parameters using fewer photons per pixel than iterative re-convolution methods.^25,26^ More recently, “smart binning” techniques based on spatiotemporal correlation between pixels^27,28^ and deep learning neural network-based algorithms^29–31^ have been developed to recover lifetimes in images with fewer than a hundred photons per pixel. However, these post-processing techniques usually incur high computational costs or require case-specific pretraining and, therefore, are not frequently used. In practice, live cell FLIM is commonly performed using commercial TCSPC electronics that are easily integrated into many imaging systems. Pixel-wise fitting (after local spatial pixel binning) using WLS or MLE^32^ is then typically performed in a graphical user interface prior to image segmentation or region of interest (ROI) selection for further statistical analysis.^13^

Here, we developed a simple and computationally cheap technique that enables fast (low photon) FLIM acquisition in a manner compatible with a widely used acquisition and analysis pipeline without the need for specialized electronics or machine learning algorithms. We first characterize the performance of “region of interest” (ROI)-binned analysis compared to conventional pixel-level analysis using a uniform fluorescence sample, investigating the effects of photon budget on accuracy and precision. We compare the performance of the ROI-binned analysis to the Cramér-Rao lower bound (CRLB) on lifetime estimation variance for the same photon budget range. We then establish the ability of ROI-binned analysis to accurately recover NAD(P)H lifetimes in living cancer cells with low photon budgets. We take advantage of the reduced photon budget requirements of the ROI-binned analysis to capture fast single cell dynamics in moving neutrophils (∼1 s acquisition per frame providing ∼1.5 photons / pixel or ∼2000 photons / cell) and beating cardiomyocytes (∼1 s / frame), and then to capture large field of view (FOV) mosaics (1.8×1.8 mm) of HeLa cells with short (1 min) acquisition times (or ∼15× larger FOV for the same acquisition time). These results demonstrate a user-friendly and computationally light approach for recovering autofluorescence lifetime parameters in biological samples. This workflow is adaptable for commonly used analysis software and preserves single cell information without the need for computationally expensive spatiotemporal correlation analysis or complex machine learning methods.

## 2. Methods

### 2.1 Theoretical lower bound of fluorescence lifetime estimation precision

The CRLB on fluorescence lifetime estimation variance was calculated for the same sample and imaging system parameters as the fluorescent solution experiment (described in Section 2.2 and 2.3) using open-source code written by Bouchet et al.^33^ and implemented in MATLAB (MathWorks, Inc.).

### 2.2 Sample preparation: fluorescence standards and live cell culture

NADH (Sigma-Aldrich 43420) was dissolved in Tris-buffered saline (diH2O, 50 mM Tris, 150 mM NaCl, pH 7.6) at a concentration of 50 µM to minimize self-quenching (that occurs above 375 µM^34^). 10 µM glucose-6-phosphate dehydrogenase (G6PDH, Sigma-Aldrich G6378) was added immediately prior to imaging to bind NADH in solution and produce multiexponential fluorescence decays (expected protein bound fraction at 10 µM is ∼ 20%^35^). A single droplet was placed on a No. 1.5 glass-bottomed imaging dish (MatTek Corporation) and imaged using a digital (scan) zoom of 10x to produce a uniformly filled FOV (30x30 µm imaged area) in the center of the scan range, as seen in Fig. 2.

PANC-1 human pancreatic (ATCC CRL-1469) and HeLa human cervical (ATCC CCL-2) adenocarcinoma cells were maintained in high-glucose Dulbecco’s modified Eagle’s medium supplemented with 10% fetal bovine serum (Gibco) and 1% penicillin/streptomycin (Gibco). 3×10^5^ cells were plated on each glass-bottomed dish 24 h prior to imaging and maintained at 37°C and 5% CO_2_.

Primary human neutrophils were isolated from the peripheral blood of a healthy donor using the MACSxpress Whole Blood Neutrophil Isolation Kit (Miltenyi Biotec), following the manufacturer’s instructions within 1 hour of blood draw. Erythrocyte depletion (Miltenyi Biotec) was performed according to the manufacturer’s instructions after isolation. The cells were resuspended in Roswell Park Memorial Institute (RPMI) 1640 Medium (Gibco) supplemented with 10% Adult Bovine Serum (ABS), and 1% penicillin-streptomycin and kept at 37°C under 5% CO_2_. For imaging, neutrophils were plated at 200,000 cells per 50 µL of medium on 35mm glass-bottom dishes (MatTek) precoated with Cell-Tak (Corning). Blood draws were performed following protocols approved by the Institutional Review Board of the University of Wisconsin–Madison (2018–0103), and informed consent was obtained from all the donors.

Cardiomyocytes were differentiated from WTC-11 human induced pluripotent stem cells following an established method.^36^ Cardiac progenitor cells were plated on Matrigel-coated glass-bottomed plates (Ibidi) on day 5 post-differentiation and maintained in RPMI (Gibco) with B27 + insulin (LifeTechnologies) at 37°C and 5% CO_2_ ^37^. Images were acquired in fully differentiated, beating cardiomyocytes (19 days post-differentiation). Cells were incubated with Mitotracker Orange CMTM (25 nM, Invitrogen) at 37°C and 5% CO_2_ for 15 min, rinsed 3 times, and immediately imaged in fresh culture medium.

### 2.3 Fluorescence lifetime imaging

Imaging was performed on an Ultima two-photon imaging system (Bruker) with an ultrafast tunable laser source (80 MHz, 100 fs pulses; Insight DS+, Spectra Physics) coupled to a Nikon Ti-E inverted microscope with TCSPC electronics (SPC-150, Becker & Hickl and Time Tagger Ultra, Swabian Instruments). NAD(P)H was excited at 750 nm and emissions collected using a 440/80 nm bandpass filter (Chroma) and GaAsP photomultiplier tube (H7422PA-40, Hamamatsu). Mitotracker Orange (cardiomyocyte mitochondrial imaging) was excited at 750 nm and emission collected using a 590/50 nm bandpass filter. The system instrument response function (IRF, full width at half maximum ∼240 ps) was acquired from the second harmonic-generated signal of urea crystals at 750 nm captured on the same detector. Samples were illuminated through a 40×/1.15 NA objective (Nikon), with ∼6 mW laser power at the sample. PANC-1 and HeLa cells were imaged with a digital zoom of 1 (300 µm FOV), neutrophils with a digital zoom of 2 (150 µm FOV), and cardiomyocytes with a digital zoom of 2.5 (120 µm FOV). Fluorescence decays (256-time bins) were acquired across 512 × 512-pixel images with a pixel dwell time of 4.8 µs. FLIM frames (*i*.*e*., single pass of the galvanometer scanner over the FOV, ∼1.5 s/frame) were saved to disk individually. The required number of consecutive FLIM frames were combined when necessary (*e*.*g*., NADH solution, PANC-1 cells), emulating FLIM images with total integration times from 1 to 120 seconds.

### 2.4 Mosaic FLIM

The Atlas Imaging feature of the Prairie View (Bruker) microscope control software was used to program a serpentine pattern of microscope stage positions covering a large area of the cell plate (1.8×1.8 mm). FLIM images were acquired sequentially and saved separately., The images were fused in post processing using the BigStitcher^38^ plugin in Fiji^39^.

### 2.5 Image denoising and ROI segmentation

Single cell NAD(P)H intensity images were denoised using convolutional neural networks (CNN) where necessary for accurate single cell segmentation. In the case of neutrophil images, a generative accumulation of photons (GAP) network^40^ designed for mitigating shot noise in photon counting datasets was trained on ∼500 noisy images of neutrophils (without the need for any noise-free ground truth images) and used to denoise the experimental sequence of 1-sec images of neutrophils before and after activation with phorbol 12-myristate 13-acetate (PMA) treatment., The pretrained CNN included in Cellpose3^41^ was used to denoise the HeLa cells mosaic intensity images. It should be noted that denoising is performed only on the intensity images for the sole purpose of generating cell masks and therefore does not change the lifetime data used in the fitting analyses. We may also use the denoised intensity image for visualization of the color-coded lifetime image.

The uniform fluorescence standard (NADH solution) images were segmented into a grid of square ROIs with 10, 20, 40, 60, 80, or 100 pixels (width) for analysis. Automated cell segmentation was performed on NAD(P)H intensity images using Cellpose^42,43^ (model ‘cyto2’ for PANC-1; custom models for neutrophils; model ‘cyto3’ for HeLa cells). The napari viewer was used to manually segment cardiomyocytes and perform any necessary mask corrections in the other cell types.

### 2.6 Pixel-wise multiexponential fitting

Pixel-wise multiexponential fitting was performed using SPCImage v 8.8 (Becker & Hickl). Two-component decays were calculated at each pixel by the following equation: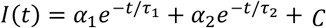, where α_1_and α_2_ are the amplitudes, τ_1_and τ_2_ are the decay lifetimes of the fast and slow decay components, respectively. *C* is the constant non-decaying background light amplitude. A binning factor of 1 (3×3 pixels) was used for all analyses.^3^ Both WLS and MLE iterative fitting and re-convolution with the IRF were used to fit multiexponential fluorescence decays.^13^

Segmentation was performed as described in 2.5 and lifetime fit parameters (α_1,_ τ_1,_ α_2_, τ_2_) were calculated for each cell or ROI using a custom Python library, ‘cell analysis tools,’^44^ that calculates the average of pixel lifetime fit parameters over the pixels of each ROI (Fig. 1).

**Fig. 1.**
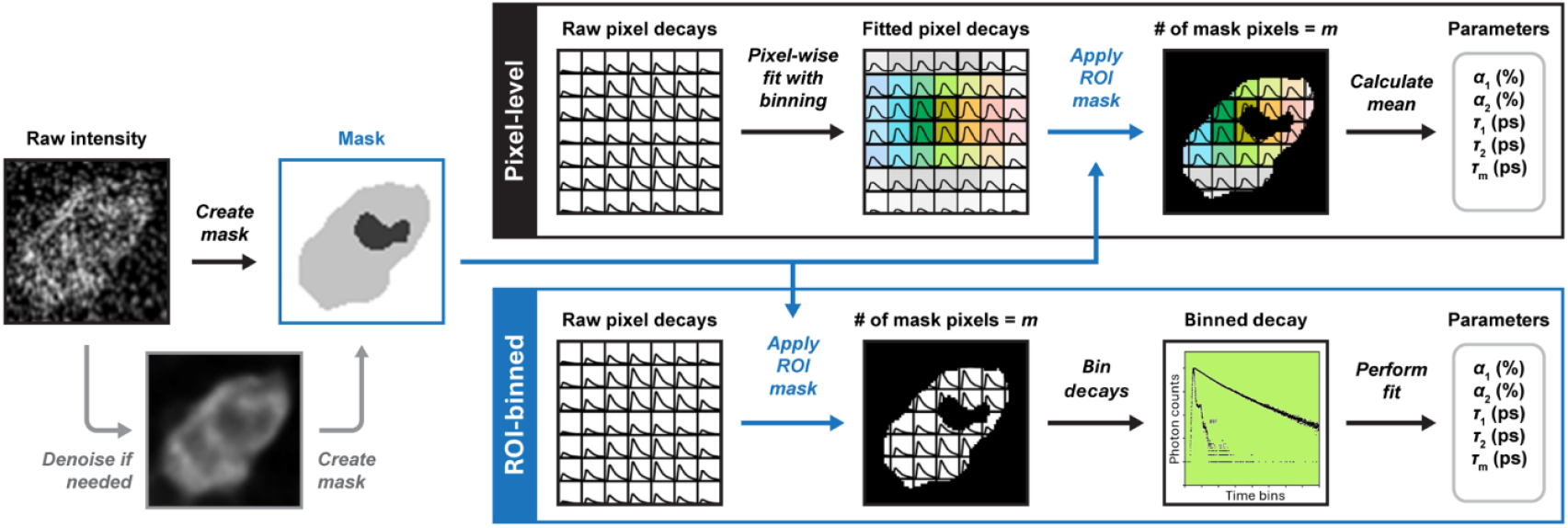
Analysis pipeline for single cell fluorescence lifetime extraction. *Pixel-level*: Raw fluorescence decays can be fitted to a multiexponential model at each pixel prior to cell segmentation. Single region/cell masks are applied, and average lifetime fit parameters are then calculated for each ROI. *ROI-binned*: Binary cell masks defining cell boundaries are applied to the raw FLIM images and a single fluorescence decay is calculated from the sum of all pixel decays contained in the ROI. Lifetime parameters are then extracted at the single-ROI level by fitting this decay.

### 2.7 ROI-binned multiexponential fitting

Segmentation was first performed on the uniform fluorescent sample or live cell images as described in 2.5 and masks were applied to the raw FLIM image files. For each mask object (*e*.*g*., single cell or ROI), the decays from all object pixels were combined to produce a decay with high photon count (Fig. 1). This aggregate “ROI-binned” decay was then identically assigned to all pixels of the object. This “new” FLIM image file was subsequently loaded and fitted in SPCImage using both WLS and MLE methods as described in 2.6. The bin factor was set to zero, since these modified images are effectively pre-binned. The lifetime fit parameters were extracted using ‘cell analysis tools.’

## 3. Results

### 3.1 NADH solution: characterizing performance with a controlled photon budget

We first characterized the performance of the pixel-level and ROI-binned methods using a uniform sample (NADH solution) and controlled photon count per pixel (Fig. 2a). The laser power was adjusted to attain a count rate of ∼1.2×10^5^ photons/s, ensuring adequate SNR while avoiding photon pile-up artifacts. We first calculated the ground-truth multiexponential lifetime parameters for the NADH solution (with G6PDH added) from the aggregation of 120 single FLIM frames. At this integration time, the average image pixel decay contains 54 photons, which is typical of autofluorescence images of cells. With sufficient pixel binning, these photon numbers can provide accurate and repeatable lifetime estimates. Both pixel-level (with a binning factor>1) and ROI-binned analysis pipelines, using either MLE or WLS fitting methods, converge on a biexponential decay profile with these parameters: *Background photons* = 10%, τ_1_= 336 ps, τ_2_ = 2371ps, α_1_= 85%, α_2_ = 15%, where background photons refers to the non-decaying ambient light intensity, also known as offset. The “ground truth” mean fluorescence lifetime (defined as τ_*m*_ = (α_1_τ_1_+α_2_τ_2_)/(α_1_+α_2_)) of NADH in solution was found to be 641 ps.

Given this ground truth, we then normalized and presented the bias and coefficient of variation (COV) across grid “cells” for lifetime estimates by each analysis pipeline (Fig. 2b-e). The independent variables in this experiment are photons per pixel on horizontal axes, and cell size, presented as the number of pixels per square grid cell.

The performance of ROI-binned analysis (orange curves) was compared to pixel-wise fitting with a bin factor of 1 (blue curves) using the MLE (Fig. 2 b-c) or the WLS (Fig. 2 d-e) methods. ROI-binned analysis results in a smaller negative bias compared to pixel-wise analysis using MLE. This indicates that pixel-wise fitting tends to underestimate lifetime, particularly at very low photons per pixel (≤ 1), a regime where ROI-binned fitting can provide usable estimates. Similarly, the COV for the ROI-binned WLS fitting pipeline is lower than its pixel-wise equivalent over the entire range of photons per pixel. However, the COV for the ROI-binned MLE fitting pipeline is only lower than the pixel-wise MLE fitting pipeline at the higher end of photons per pixel (where the pixel-wise fitting bias has shrunk to <5%).

Given the previous results, we compared the standard deviation of our unbiased lifetime estimators (MLE, WLS) to the theoretical minimum value given by the Cramér-Rao lower bound (Fig. 2f).The CRLB is presented for the mono-exponential background-free decay case (*i.e.* the shot noise limit 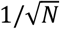), mono-exponential decay with background (3 estimated parameters), and biexponential decay with background (5 estimated parameters) with the latter being applicable to the NADH solution. The theoretical curves were calculated using the ground truth solution lifetime parameters and a cell size of m = 400 pixels. The ROI-binned standard deviation approaches the CRLB, with the ROI-binned MLE fitting being closer to the lower bound than the WLS fitting. In summary, ROI-binned fitting lowers both estimation bias and COV compared to pixel-wise fitting by combining decay photons belonging to the same cell object prior to performing the fit, thus alleviating the effects of photon shot noise more effectively.

### 3.2 Comparison of pipelines in live cell images at varying integration times

Having characterized the performance of each pipeline in a uniform solution, we next performed a real-world comparison of the pipelines in FLIM images of cancer cells. In addition to pixel-wise and ROI-binned fitting pipelines, we also processed the images using CASPI,^27^ a computational pre-processing algorithm that collaboratively uses local and non-local temporal correlations between pixel decays to recover decay profiles in low-light regimes. At the cost of additional computation, the CASPI algorithm effectively spatially and temporarily denoises the 3D FLIM data cube while preserving pixel-level spatial resolution.

A single FOV of PANC-1 cells was imaged using multiphoton FLIM with varying integration times (1 to 120 s) as before. The photon counts were maintained at a constant rate (2×10^5^ photons/s). The FLIM images were analyzed one of three ways: conventional pixel-wise MLE fitting with a binning factor of 1 (3×3 pixels), ROI-binned MLE fitting, or pixel-wise MLE fitting with a binning factor of 1 after pre-processing using the CASPI algorithm^27^ before extracting single cell lifetime parameters (Fig. 3). The ground-truth NAD(P)H mean lifetime (τ_*m*_) was calculated using pixel-wise MLE fitting of 120 s acquisition, and mean lifetimes converged to the true value (<5% bias) after ∼15 s.

**Fig. 2.**
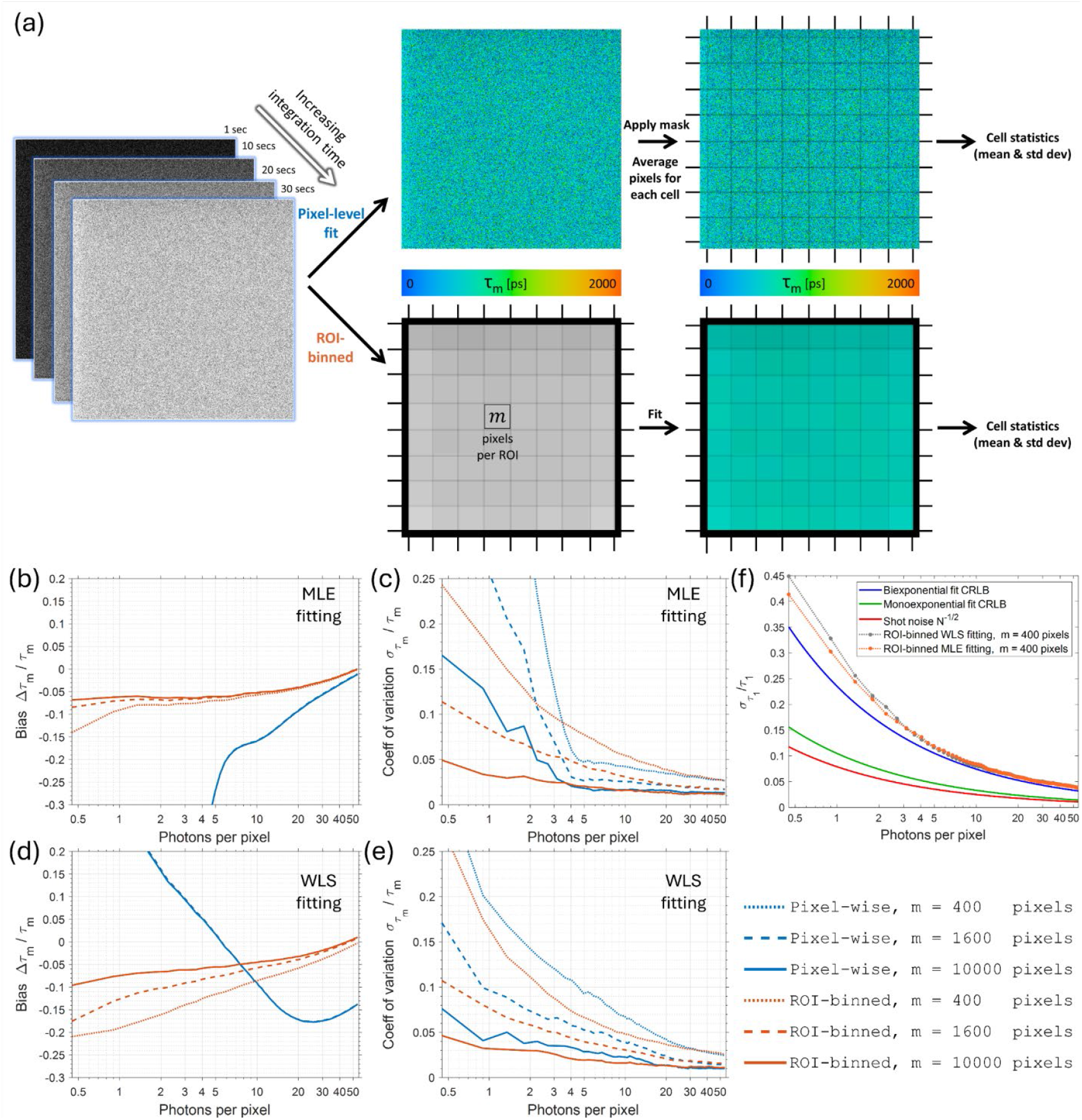
Uniform NADH solution phantom characterizes performance of fitting methods. (a) A uniform solution of 50 µM NADH + 10 µM G6PDH was imaged as a sequence of 120 single FLIM frames (∼1 sec each) that were combined in post-processing to yield integration times ranging from 1 to 120 seconds (equivalent to 0.45 to 54 photons / pixel). The resulting FLIM images were then divided into a grid of squares simulating single cells (several grid sizes) and analyzed using pixel-wise or ROI-binned fitting methods in SPCImage. (b,c) The normalized bias and standard deviation of the estimates using MLE fitting. (d,e) The normalized bias and standard deviation of the estimates using WLS fitting. (f) The normalized standard deviation of the τ_1_ lifetime estimates from the ROI-binned MLE and WLS fitting analyses approach to the theoretical CRLB calculated using the “ground truth” lifetime of the NADH solution.

**Fig. 3.**
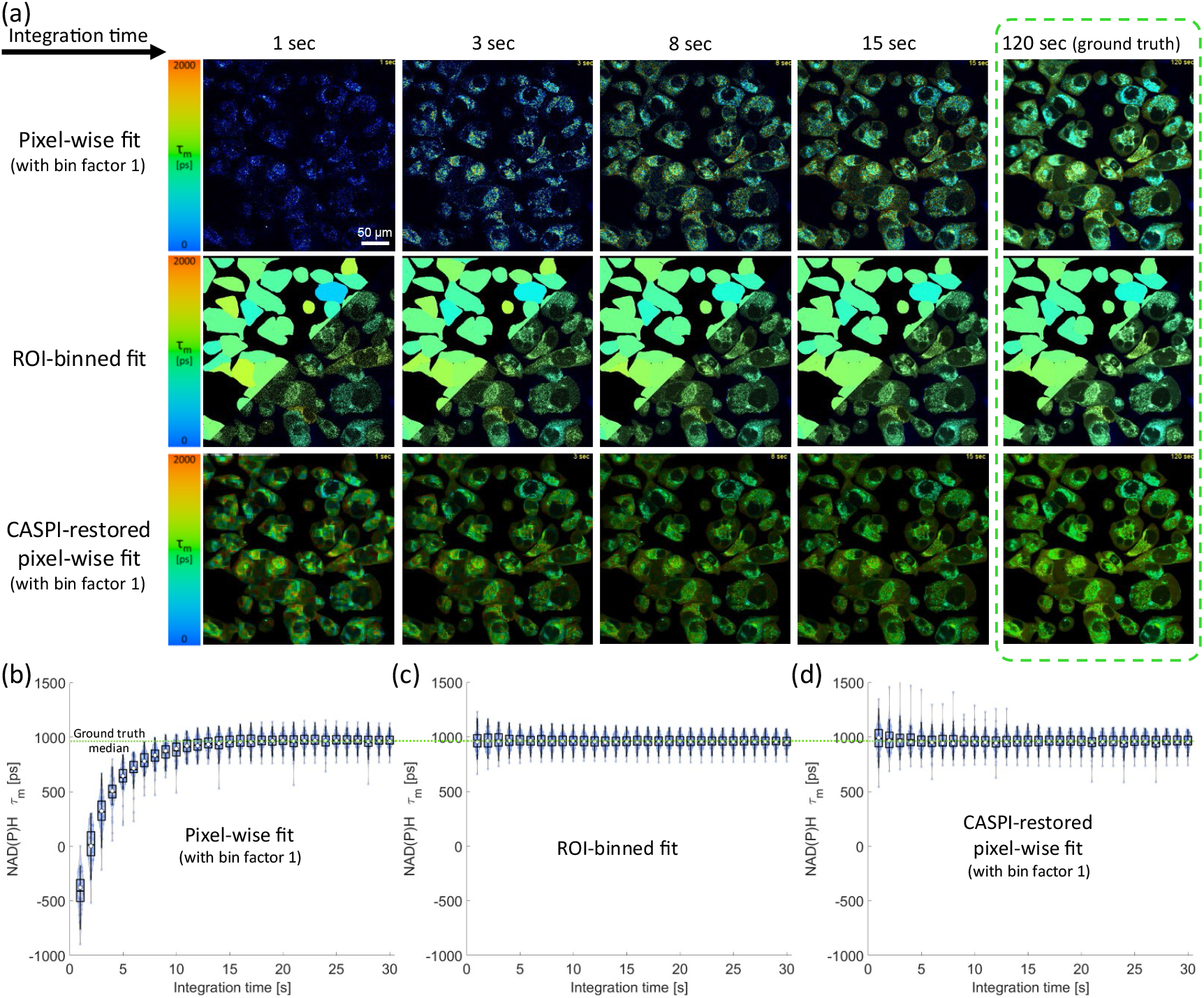
ROI-binned analysis produces valid autofluorescence lifetime estimates within one second of integration. Adherent PANC-1 cells were imaged as a real-world test case. (a) Representative FLIM images with increasing integration times when analyzed using the conventional pixel-wise MLE fitting, ROI-binned MLE fitting, or CASPI restoration followed by pixel-wise MLE fitting, all using SPCImage software. Violin plots of the single cell mean lifetimes distribution with increasing integration times are given for the (b) conventional pixel-wise MLE fitting, (c) ROI-binned MLE fitting, and (d) CASPI-restored pixel-wise MLE fitting. The ROI-binned and CASPI-restored FLIM images provide valid lifetime estimates from the first second of integration, but conventional pixel-wise fitting only converges to the true lifetimes after about 15 s of integration.

Pixel-wise MLE fitting produces invalid negative lifetime estimates in the first 3 seconds of integration due to very low photons per pixel (Fig. 3b). In contrast, ROI-binned MLE analysis produces estimates within 5% bias with just 1 s integration (Fig. 3c). The resulting lifetime images are presented with flat lifetime color-coding alone, or superimposed on pixel-level intensity for visualization (Fig. 3a). Like ROI-binned MLE analysis, CASPI restoration before fit analysis results in pixel-level estimates within 5% bias with just 1 s integration (Fig. 3d). Video S1 presents FLIM images produced by these pipelines for the first 15 s of integration.

### 3.3 Application: dynamic imaging of primary neutrophils

Neutrophils are highly dynamic and fast-responding innate immune cells. Conventional FLIM with minute-long integration times fails to capture rapid metabolic shifts in neutrophils, and the movement of neutrophils leads to blurred images (Fig. 4b). To test the feasibility of ROI-binned analysis for capturing these dynamic processes, we applied the pipeline to single-frame (∼1 s each) FLIM images of these neutrophils immediately before and after activation with PMA. To this end, single-frame cell masks are required to identify pixels corresponding to individual cells. However, attempting to segment these single-frame intensity images in Cellpose^43^ leads to failure due to the effects of shot noise at these low photon counts (Fig. 4 c). We trained and applied a GAP denoising neural network^40^ to mitigate the shot noise in these single-frame intensity images. As a result of this denoising, Cellpose segmentation of single cells in single-frame images is now successfully performed (Fig. 4 d).

**Fig. 4.**
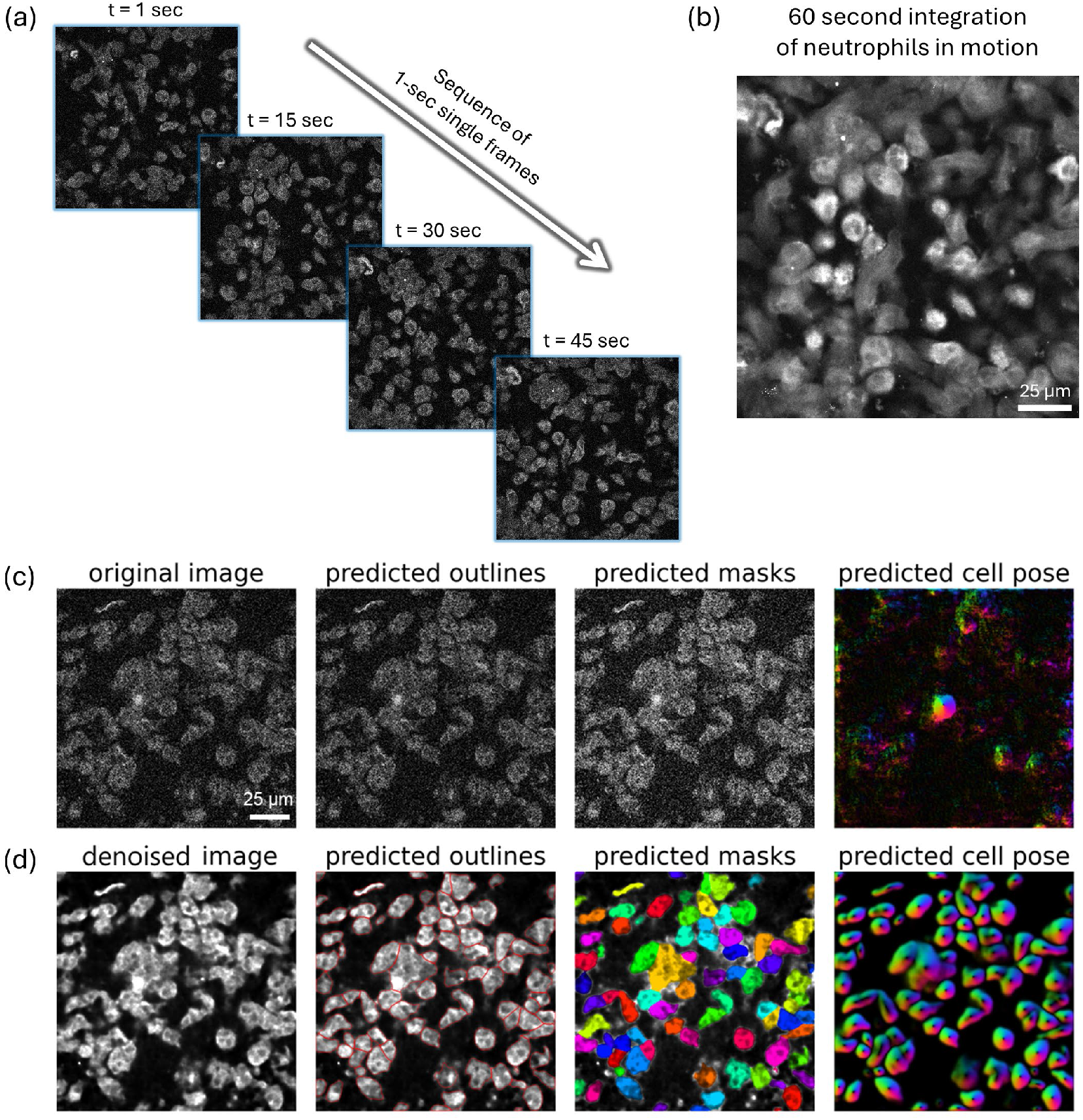
CNN-based denoising enables single cell segmentation and lifetime estimation for fast FLIM imaging. (a) Representative intensity images of migrating neutrophils captured as a series of single-frame FLIM images. (b) Combining all frames for a conventional 60 s integration time results in a blurred image that cannot be segmented and FLIM analysis would suffer from crosstalk between cells. FLIM images would ideally be analyzed on a per-frame basis. However, cell segmentation in Cellpose fails (c) due to heavy influence of shot noise in single-frame intensity images. (d) Denoising single-frame images using a GAP neural network enables successful single cell segmentation necessary for ROI-binned analysis.

Representative raw, denoised, and ROI-binned mean lifetime images of the neutrophils before and one minute after activation with PMA are shown in Fig. 5a. The single cell mean lifetime distributions over the course of six minutes (1 min before and 5 min after PMA activation) are plotted in Fig. 5b in 1 s intervals. NAD(P)H mean lifetimes are stable over one minute in control neutrophils. In comparison, upon treatment with PMA, the mean lifetime drops within the first minute and subsequently begins to slowly recover over the next few minutes (Video S2). These rapid changes would not be visible with the much longer acquisition time typically required for autofluorescence FLIM, showcasing a clear use for this analysis method.

**Fig. 5.**
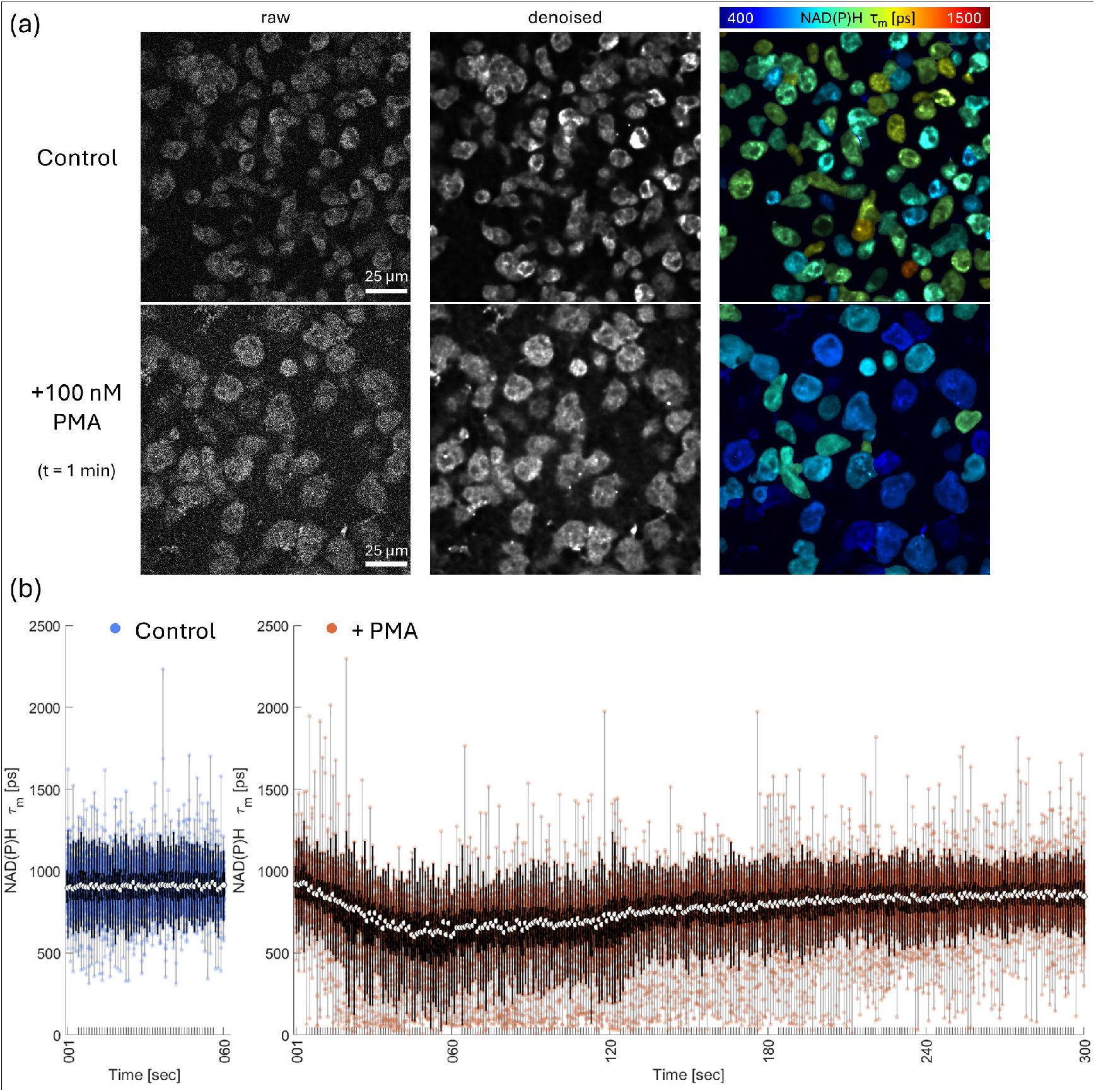
ROI-binned FLIM imaging captures fast metabolic dynamics in activated migrating neutrophils. (a) Representative raw intensity, denoised intensity, and ROI-summed mean lifetime superimposed on denoised intensity images of neutrophils before and one minute after treatment with PMA. (b) Time series boxplots of single cell NAD(P)H lifetime distributions in the same FOV for 60 seconds before (left, control, blue) and 5 minutes after (right, +PMA, red) treatment with PMA in one-second intervals. This 1 s resolution reveals fast metabolic dynamics with the mean lifetime dropping for one minute after PMI treatment and gradually recovering over the next few minutes. White dots represent the median value, dark box shows the interquartile range, and whiskers indicate ±1.5× interquartile range..

### 3.4 Application: Mitochondrial imaging in beating cardiomyocytes

We next used the ROI binning method for recovering NAD(P)H lifetimes in conjunction with organelle staining to visualize dynamic, subcellular lifetime information in beating cardiomyocytes. These cells are highly metabolically active and known to have extensive mitochondrial networks. A sequence of 60 single-frame fluorescence lifetime images was acquired in two spectral channels, capturing NAD(P)H autofluorescence and MitoTracker Orange simultaneously. An Otsu thresholding of the MitoTracker intensity channel was used to generate mitochondrial network masks which were then combined with single whole cell masks to produce single cell mitochondrial network ROIs. The NAD(P)H fluorescence lifetime decays were ROI-binned and analyzed within these single cell mitochondrial networks. Mean lifetime values for mitochondria were color-coded and presented as seen in Video S3. Mitochondrial mean fluorescence lifetime appears to change rapidly throughout different phases of the beat cycle and is heterogeneous across cells. This example demonstrates that the ROI binning pipeline can be combined with subcellular organelle labels to perform fast autofluorescence FLIM of specific cell structures.

### 3.5 Application: Mosaic imaging of large FOV

Autofluorescence FLIM of large FOVs is often time-consuming owing to long integration time requirements. ROI-binned lifetime analysis would considerably increase the image acquisition speed in such cases, enabling stitching of many FOVs to cover a large area. Mosaic images consisting of 6×6 FOV tiles (1.8×1.8 mm^2^) were acquired of HeLa cells using a serpentine pattern of stage locations on the multiphoton microscope. A ground truth mosaic was acquired by integrating the FLIM image at each position for 15 passes of the scanning galvos (∼22 s), requiring a total acquisition time of about 14 minutes. A fast mosaic was then acquired by taking a single FLIM frame (∼1.5 s) at each position for a total acquisition time < one minute. In post-processing, the images were fused using the BigStitcher^38^ plugin in Fiji^39^ software and segmented in Cellpose3 using the built-in pretrained one-click denoising neural network^41^. Both datasets were analyzed using ROI-binned MLE fitting and color-coded lifetime values were superimposed on denoised intensity images as shown in Fig. 6a. We compared the single-cell NAD(P)H mean lifetime (τ_m_, Fig. 6c) and free fraction (α_1_, Fig. 6e) measured in the ground truth (15 frames integrated) and fast FLIM (single frame) mosaics and present the difference between the lifetime variables measured from each mosaic (Fig. 6d, f). The FLIM images are visually similar, and the quantified difference suggests that taking a single FLIM frame of each FOV only results in a -1.6% mean lifetime (τ_m_) bias or a -0.1% proportion of short lifetime (α_1_) bias in extracted single cell lifetime parameters while reducing total imaging time by a factor of ∼14.

**Fig. 6.**
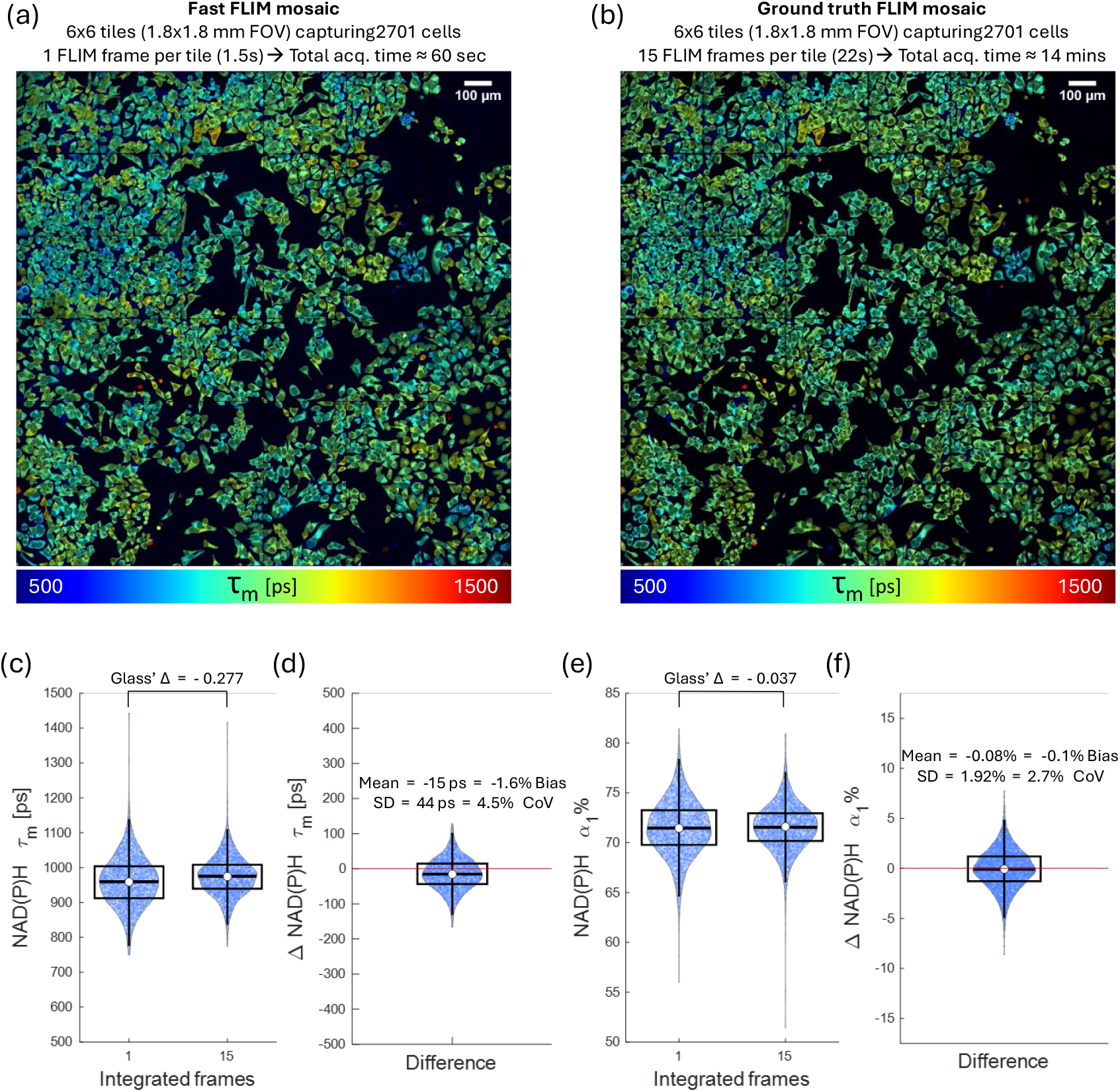
Fast large-FOV mosaic FLIM imaging using ROI-binned analysis. (a) 6×6 fast FLIM mosaic using a single frame (∼1.5 s/frame) at each tile position captured 2701 HeLa cells within one minute. In contrast, acquiring the (b) “ground truth” FLIM mosaic by accumulating 15 frames at each tile position requires 14 minutes of total integration time. Both mosaics were analyzed using ROI-binned MLE fitting and the color-coded lifetime values are superimposed on the corresponding denoised intensity image. (c) Distribution of single-cell NAD(P)H mean lifetimes for single-frame and 15-frame integration. (d) Distribution of mean lifetime difference between the two FLIM images, showing an average bias of -1.6%. (e) Distribution of single cell free fraction of NAD(P)H for single-frame and 15-frame integration. (f) Distribution of free fraction difference between the two FLIM images showing an average bias of -0.1%. Box plot shows the interquartile range and whiskers extend from the box to 1.5× the interquartile range. White dot shows the median, and horizontal line the mean.

## 4. Discussion

We presented an ROI-binned lifetime fitting method as a photon-efficient alternative to pixel-wise fitting that recovers accurate single-cell lifetime parameters. Along with a considerable decrease in FLIM acquisition times, this approach results in improved accuracy (lower bias) and precision (lower standard deviation) of lifetime estimates compared to pixel-wise fitting with the same photons per pixel. We experimentally quantified this method using a uniform NADH solution for a complex decay fitting scenario with realistic photon counts and instrument effects. This approach is a natural extension of spatial pixel binning, which is already common practice for increasing photon counts during pixel-wise fitting analysis. However, we aggregated all pixels in the same cell or ROI in the image rather than blindly grouping neighboring pixels (3×3, 5×5, *etc*.), as is common with typical spatial binning. We further demonstrated these advantages in a real-world live cell imaging case by analyzing FLIM images from PANC-1 cells acquired over a range of photons per pixel.

The increased photons per decay afforded by ROI-binning enabled us to image both faster single cell dynamics and larger mosaic FOVs compared to pixel-wise fitting. We demonstrated the ability to track fast metabolic dynamics in motile, activated neutrophils and beating cardiomyocytes with FLIM frame rates of about 1 frame per second. Machine learning advancements in image denoising under shot noise conditions (which is the dominant noise in TCSPC FLIM) enabled successful segmentation of single cells in noisy fast FLIM images which, in turn, enabled ROI-binning for successful single-cell lifetime estimation.

This ROI-binned fitting approach is hardware agnostic and can be applied to TCSPC images from any multiphoton or confocal system to improve estimation accuracy, increase acquisition speed, and reduce computational cost (since fewer iterative fitting operations are needed compaerd to pixel-wise fitting). The ROI can be defined as any desired group of pixels for either whole cell binning (e.g., PANC-1, HeLa, neutrophils) or specific subcellular regions (e.g., mitochondrial network in cardiomyocytes). Thus, ROI can be defined as cytoplasm, nucleus, mitochondria, or other organelles with or without the assistance of live cell labels. Irrespective of the ROI, binning pixels together prior to re-convolution fitting results in improved fitting performance compared to pixel-wise fitting followed by averaging.

In situations where preserving pixel-level spatial resolution is desired, global fitting, smart binning algorithms like CASPI, or other machine learning algorithms that restore pixel decays under low photon conditions or directly recover lifetime parameters in fit-free fashion can be used^23,27,45,46,28,47^. However, these techniques either have increased computational cost or require the algorithm to be pre-trained for the sample of interest.

In conclusion, ROI-binned analysis is a photon-efficient approach to estimate single-cell fluorescence lifetime parameters that, depending on the user priorities, can improve estimation accuracy and precision, or increase imaging speed by a factor of 15 or more. ROI-binned analysis enables autofluorescence FLIM of fast dynamics that would otherwise not be achievable using laser-scanning FLIM, all while lowering the computational cost of single cell analysis.

## Supporting information

Supplemental Video 1 - PANC1

Supplemental Video 2 - Neutrophils

Supplemental Video 3 - Cardiomyocytes

## Acknowledgement

The authors thank Professor Mohit Gupta for early sharing of the source code for the CASPI algorithm.

## Code and Data Availability

The source code is available upon request and will be released as open source upon publication. An example dataset of PANC-1 cells and basic ROI binning source code are available at https://github.com/skalalab/ROI_binning_bioarchive

## Abbreviations

CNN: convolutional neural network
COV: coefficient of variation
CRLB: Cramér-Rao lower bound
FAD: flavin adenine dinucleotide
FLIM: fluorescence lifetime imaging
FOV: field of view, GAP: generative accumulation of photons
IRF: instrument response function
MLE: maximum-likelihood estimation
NAD(P)H: nicotinamide dinucleotide adenine (phosphate)
PMA: phorbol 12-myristate 13-acetate
ROI: region of interest
SNR: signal-to-noise ratio
SPAD: single photon avalanche diode
TCSPC: time-correlated single photon counting
WLS: weighted least squares

